# *Tc*HRG plays a central role in orchestrating heme uptake in *Trypanosoma cruzi* epimastigotes

**DOI:** 10.1101/2023.04.05.535753

**Authors:** Evelyn Tevere, Cecilia Beatriz Di Capua, Nathan Michael Chasen, Ronald Drew Etheridge, Julia Alejandra Cricco

**Author notes:** **CORRESPONDING AUTHOR:** Julia A. Cricco, **E-mail:**.

## Abstract

*Trypanosoma cruzi*, a heme auxotrophic parasite, can control intracellular heme content by modulating *Tc*HRG expression when a free heme source is added to axenic culture. Herein, we explore the role of *Tc*HRG protein in regulating the uptake of heme derived from hemoglobin in epimastigotes. It was found that the parasite’s endogenous *Tc*HRG (protein and mRNA) responds similarly to bound (hemoglobin) and free (hemin) heme. Additionally, the overexpression of *Tc*HRG leads to an increase in intracellular heme content. The localization of *Tc*HRG is also not affected in parasites supplemented with hemoglobin as the sole heme source. Endocytic null epimastigotes do not show a significant difference in growth profile, intracellular heme content and *Tc*HRG protein accumulation compared to WT when feeding with hemoglobin or hemin as a source of heme. These results suggest that the uptake of hemoglobin-derived heme likely occurs through extracellular proteolysis of hemoglobin *via* the flagellar pocket, and this process is governed by *Tc*HRG. In sum, *T. cruzi* epimastigotes controls heme homeostasis by modulating *Tc*HRG expression independently of the source of available heme.

## INTRODUCTION

*Trypanosoma cruzi* is the causative agent of Chagas disease, which is a widespread parasitic disease in the Americas [1]. This parasite undergoes a complex life cycle, which involves a mammalian host and a triatomine insect vector. *T. cruzi*, like other trypanosomatids that cause neglected diseases in humans such as *Trypanosoma brucei* and *Leishmania spp*., are heme auxotrophic [2,3] and rely on obtaining this essential cofactor from their hosts or vectors. In the triatomine midgut, hemoglobin (Hb) derived from the bloodmeal is subjected to proteolysis, leading to the release of the heme moiety. Therefore, *T. cruzi* epimastigotes in their natural habitat encounter both Hb bound and free heme.

*In vitro* studies have shown that epimastigotes are able to incorporate both free and Hb derived heme to supply its metabolic requirements. Evidence suggests that, although internalized following different kinetics or pathways, both molecules are ultimately stored in reservosomes [4] which are lysosome-related organelles only present *T. cruzi* epimastigotes. It has been established that trans-membrane proteins belonging to the Heme Response Gene (HRG) family [5] are involved in the transport of heme from the environment in trypanosomatids. Several HRG family members have been described in trypanosomatids, for example LHR1 (*Leishmania Heme Response 1*) in *Leishmania amazonensis* [6–8], *Tb*HRG (*Trypanosoma brucei Heme Responsive Gene*) in *T. brucei* [9,10], and *Tc*HRG (previously named *Tc*HTE, *Trypanosoma cruzi Heme Transport Enhancer*) in *T. cruzi* [11,12]. Recently we reported a direct relationship between *Tc*HRG and heme uptake based on the expression profile of the endogenous *Tc*HRG gene and intracellular heme status. *Tc*HRG (mRNA and protein) is mainly detected in the replicative life cycle stages of *T. cruzi* (epimastigote and amastigote), in which heme uptake is observed [11]. Also, *Tc*HRG is highly expressed when epimastigotes are incubated with low or scarce levels of heme and becomes undetectable when intracellular heme reaches an optimal range. This suggests that epimastigotes can sense intracellular heme and modulate *Tc*HRG expression to control free heme uptake [12].

On the other hand, Hb uptake *via* receptor-mediated endocytosis at the flagellar pocket (FP) was demonstrated in *T. brucei* [13] and *Leishmania spp*. [14,15], but these endocytic phenomenon through the FP have not been observed in *T. cruzi*. Moreover, it has not been found any specific cargo receptor in this parasite. Instead, *T. cruzi* has a specialized organelle called the cyto<u>s</u>tome-cyto<u>p</u>harinx <u>c</u>omplex (SPC) which is involved nutrient acquisition. The SPC is an invagination of the plasma membrane at the anterior end of the cell, which penetrates the cytoplasm towards the posterior end of the cell [16]. In the epimastigote stage, endocytosed proteins and lipids enters the cell through the cytostome, are transported *via* endolysosomal vesicles through the cytopharynx and are finally stored in reservosomes at the posterior end of the cell [17]. The SPC and the reservosomes are absent in *T. brucei* and *Leishmania spp*, constituting other relevant differences between them and *T. cruzi*.

In this report, we explored the utilization of Hb as a heme source in epimastigotes of *T. cruzi* and examined the role of *Tc*HRG in Hb-derived heme homeostasis by analyzing endogenous *Tc*HRG expression, the effect of the overexpression of r*Tc*HRG, and the abolition of endocytosis when cultured in Hb-supplemented medium. We show that endogenous *Tc*HRG responded similarly to Hb as it did to free heme (added as hemin) at both the mRNA and protein level. Also, the intracellular heme content was increased in parasites that overexpress recombinant *Tc*HRG and was not affected in endocytic-null parasites when Hb was used as a heme source. Our results support an extended model for heme homeostasis in *T. cruzi* epimastigotes that includes both heme sources. We postulate that “free heme” obtained after extracellular Hb degradation is the main pathway for Hb-derived heme uptake in epimastigotes and it is enhanced and controlled by the heme transporter *Tc*HRG.

## RESULTS

### *Tc*HRG responds to hemoglobin

To study the effect of Hb as heme source, we followed the growth profile of WT epimastigotes comparing both free heme (added as hemin) and Hb-derived heme. Briefly, parasites cultured in LIT-10% FBS + 5 μM hemin [12] were challenged to heme starvation for 48 h and then transferred to media supplemented with 5, 20, and 50 μM hemin, with 1.25, 5, and 12.5 µM Hb (equivalent to 5, 20, and 50 µM heme as Hb) and without any source of heme added (0 μM). The number of parasites per ml was measured every day for 14 days, performing a dilution in fresh media on the seventh day. Along the 14 days, the growth of Hb-supplemented epimastigotes was slightly inferior compared to the standard condition (LIT-10% FBS + 5 μM hemin). During the second week, Hb-supplemented epimastigotes displayed similar growth profiles among each other, in stark contrast with the negative effect observed on epimastigotes’
s growth at higher hemin concentrations, 20 and 50 μM. (Fig. 1A and [12]). We did not detect any alteration in parasite morphology when the Hb-supplemented epimastigotes were observed under an optical microscope, contrary to the alterations caused by equivalent free heme concentrations added as hemin [12].

**Figure 1:**
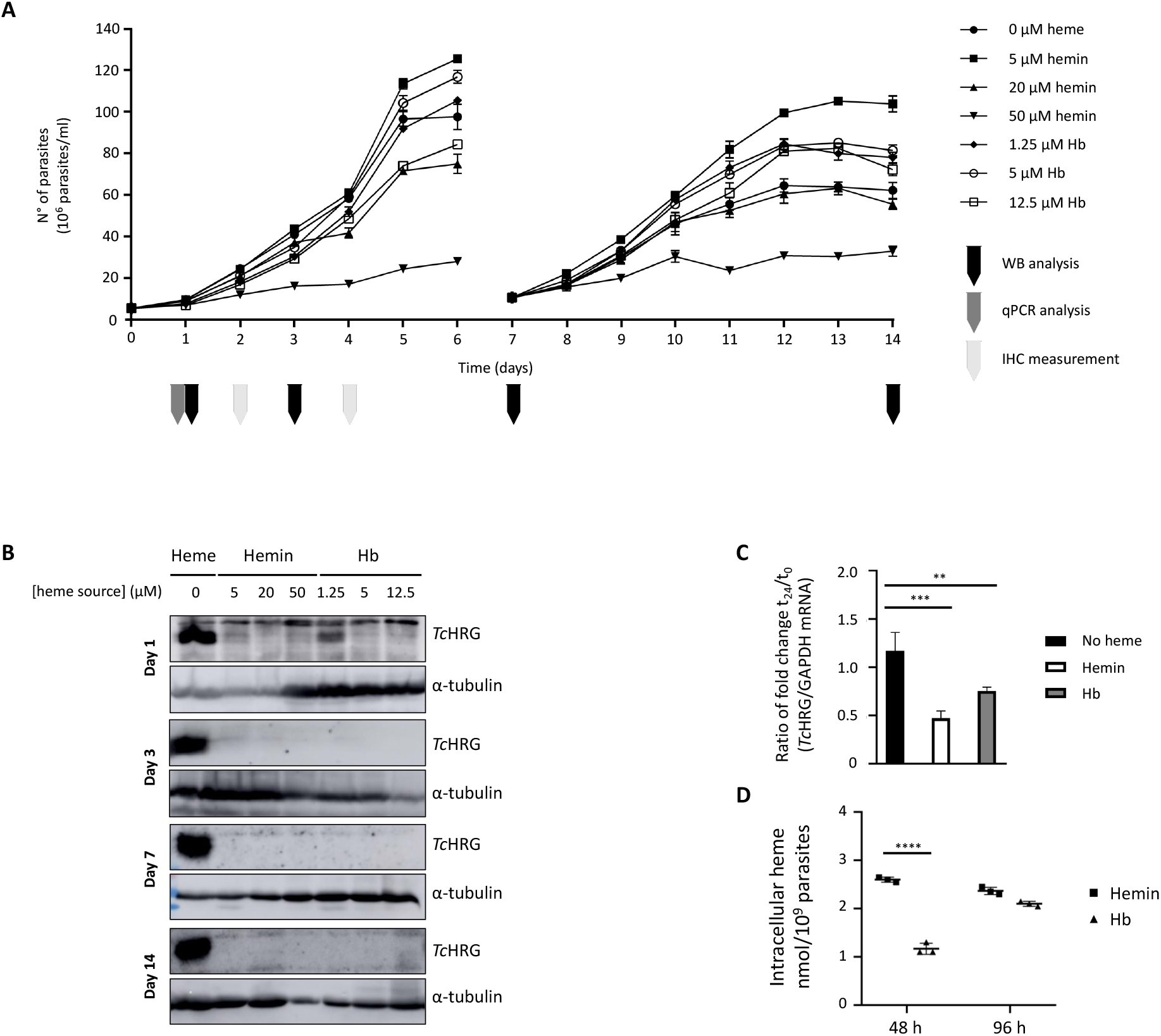
Endogenous *Tc*HRG expression responds to Hb-derived heme in epimastigotes. WT epimastigotes (Dm28c) were routinely cultured in LIT-10% FBS + 5 µM hemin, then challenged to heme starvation for 48 h. After heme starvation, epimastigotes were collected, washed with PBS and transferred to culture media were supplemented with 5, 20, or 50 µM hemin, 1.25, 5, and 12.5 µM Hb, or without heme (0 µM). (A) Growth curve of WT epimastigotes supplemented with Hb, with hemin or without the addition of heme source for 14 days. The number of cells was followed daily for 7 days. A dilution to the initial concentration in fresh medium as performed at day 7, and the growth curves were followed for another 7 days. Data are presented as mean ± SD of three independent assays. Microcentrifuge tubes cartoons in different colors indicate sampling for Western blot (WB) analysis (black), quantitative PCR (qPCR) analysis (dark gray) and intracellular heme content (IHC) measurements (light gray) on the corresponding days. (B) Detection of endogenous *Tc*HRG by Western blot. Samples were taken on days 1, 3, 7, and 14 over the course of the growth curves. Polyclonal anti-*Tc*HRG antibodies were used to recognize endogenous protein in total extracts of epimastigotes. Detection of α-tubulin was used as a loading control. (C) Quantification of *Tc*HRG mRNA levels in WT epimastigotes cultured in media with 5 µM hemin, 1.25 µM Hb or without heme source. Samples were taken at t_0_ (after heme starvation and prior incubating parasites in the different conditions) and t_24_ (24 h post treatment). mRNA was quantified by qRT-PCR. GAPDH was used for normalization. Data are presented as mean ± SD of three technical replica, expressed as the ratio of fold change of t_24_ to t_0_. Statistical significance was determined by one-way ANOVA followed by Dunnett’s multiple comparisons test (***, p < 0.001; **, p < 0.01). (D) IHC determined by pyridine method of epimastigotes incubated in media with 5 µM hemin or 1.25 µM Hb. Samples were taken after 48 and 96 h of treatment. Data are presented as mean ± SD of 3 independent assays. Statistical significance was determined by two-way ANOVA followed by Sidak’s multiple comparisons test (****, p < 0.0001).

We then evaluated *Tc*HRG protein accumulation by Western blotting on samples taken after one-, three-, seven-, and fourteen-days post addition of a heme source. To recognize the endogenous protein, anti-*Tc*HRG polyclonal antibodies were used (named anti-*Tc*HTE in[12]). Fig. 1B shows that *Tc*HRG corresponding signal was detected as an intense band in parasites incubated without heme throughout the assay. Also, a weak signal was observed in parasites incubated with 1.25 μM Hb on day 1. *Tc*HRG was almost undetectable since day 1 in the remaining conditions.

In addition, *Tc*HRG expression was also examined through qRT-PCR analysis 24 h upon treatment with 5 µM hemin or 1.25 µM Hb (Fig. 1C). Consistently, *Tc*HRG mRNA amounts remained constant in heme-starved epimastigotes. Likewise, significant reductions of approximately 50% and 25% in mRNA levels were observed 24 h after treatment with 5 μM heme as hemin and Hb, respectively.

On the other hand, the intracellular heme content (IHC) of parasites supplemented with 5 µM hemin or 1.25 µM Hb for 48 and 96 h was analyzed. Fig. 1D shows that, after 48 h of treatment, parasites incubated with hemin reached 2.6 ± 0.1 nmol heme/10^9^ parasites, meanwhile parasites incubated with Hb showed a significant lower amount of IHC, about 1.2 ± 0.1 nmol heme/10^9^ parasites, approximately 50%. After 96 h, parasites incubated with hemin maintained IHC and those incubated with Hb increased their IHC to 2.1 ± < 0.1 nmol heme/10^9^ parasites, similar to hemin treated ones.

In summary, epimastigotes tolerated higher heme concentrations when Hb was the source. They reached similar IHC when incubated with equivalent concentrations of heme using both sources, presumably the optimal intracellular heme content under these experimental conditions. However, epimastigotes incubated with Hb required more time to reach IHC than those incubated with hemin. Additionally, endogenous *Tc*HRG (protein and mRNA) accumulation generated by heme starvation underwent a significant decrease in response to the addition of Hb.

### Endocytic null parasites maintain heme homeostasis

To gain insight in the uptake of Hb-derived heme, we evaluated what happened in epimastigotes unable to perform endocytosis when incubated with Hb as a heme source. We used Δ*Tc*Act2 epimastigotes from Y strain, in which the gene for the actin isoform 2, which is necessary for SPC-mediated endocytosis, was deleted (manuscript in preparation). The WT line and a line in which the *Tc*Act2 gene was added back (complemented) with a Ty tag (*Tc*Act2.Ty) were used as controls.

First, to evaluate the behavior of the endocytic null parasites, we registered the growth performance of the three cell lines (after 48 of heme starvation treatment) in a medium supplemented with 5 µM hemin, 1.25 µM Hb, or without the addition of any source of heme (0 μM) for seven days. As shown in Figure 2A, the growth of the Δ*Tc*Act2 was comparable in both heme sources, although the maximal parasite/ml number was slightly less when compared to the WT and *Tc*Act2.Ty lines in same conditions. Additionally, the growth of the three lines was severely impaired when no heme source was added. Endogenous *Tc*HRG expression was also evaluated by Western blotting after 48 h of incubation in the conditions mentioned above. As shown in Figure 2B, *Tc*HRG signal in WT, Δ*Tc*Act2, and *Tc*Act2.Ty was clearly detected in samples incubated without the addition of heme and in Hb-supplemented ones. Also, a very weak signal was observed in WT, Δ*Tc*Act2, and *Tc*Act2.Ty parasites supplemented with hemin.

**Figure 2:**
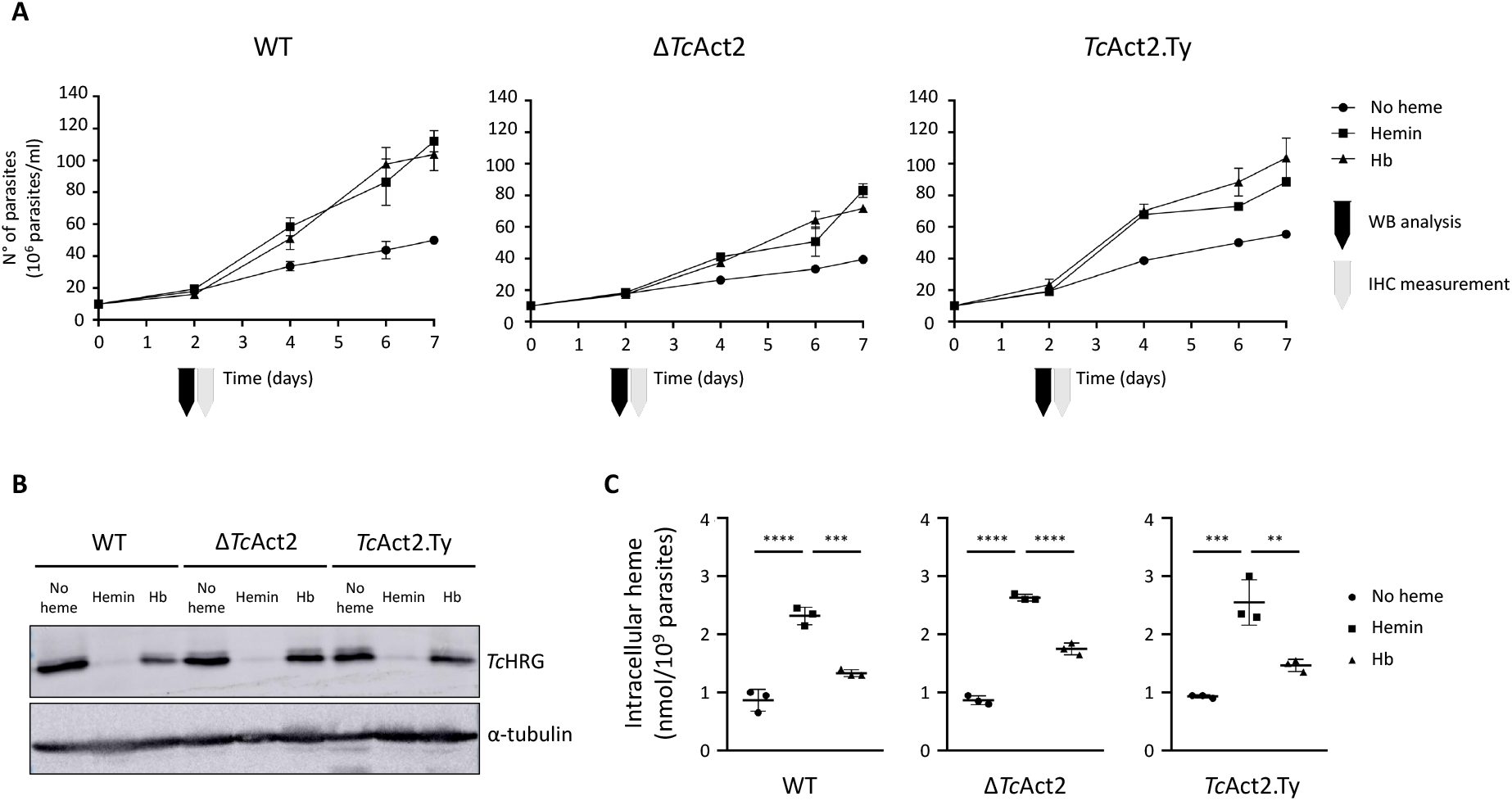
Endocytic null epimastigotes are able to grow in Hb-supplemented medium and to reach WT intracellular heme values. WT, *Tc*Act2 and *Tc*Act2.Ty (Y strain) epimastigotes were routinely cultured in LIT-10% FBS + 5 µM hemin, subjected to heme starvation for 48 h, as described previously, and finally transferred to media supplemented with 5 µM hemin, 1.25 µM Hb, or without heme source added (0 µM) for one week. After 48 h, samples were taken to perform Western blot analysis and intracellular heme measurements. Microcentrifuge tubes cartoons indicate sampling for Western blot (WB) analysis (black), and intracellular heme content (IHC) measurements (light gray) on the corresponding days. (A) Growth curve of WT, *Tc*Act2 and *Tc*Act2.Ty epimastigotes cultured in media 5 µM hemin, 1.25 µM Hb, or without heme added. The number of parasites was followed daily for one week. Data are presented as mean ± SD of three independent assays. (B) Western blot assay using anti-*Tc*HRG antibodies to detect endogenous *Tc*HRG protein in total extracts of WT, *Tc*Act2 and *Tc*Act2.Ty epimastigotes incubated for 48 h in media supplemented with 5 µM hemin, 1.25 µM Hb or without heme source. α-tubulin was used as a loading control. (C) IHC determined by pyridine method of WT, *Tc*Act2 and *Tc*Act2.Ty epimastigotes incubated for 48 h in media supplemented with 5 µM hemin, 1.25 µM Hb or without heme source. Data are presented as mean ± SD of 3 independent assays. Statistical significance was determined by one-way ANOVA followed by Dunnett’s multiple comparisons test. (****, p < 0.0001; ***, p < 0.001; **, p < 0.01).

Additionally, we analyzed the IHC of these parasites after 48 h of incubation with both heme sources and without heme. Figure 2C shows that IHC profiles were similar in the three lines. All of them reached an IHC of about 2.3-2.6 nmol heme/10^9^ parasites in hemin-supplemented media, while when they were incubated with Hb the IHC was significantly lower, about 1.3-1.8 nmol heme/10^9^ parasites. Also, when no heme source was added (0 µM), the three variants presented IHC that were significantly lower as compared to parasites treated with hemin, about of 0.9 nmol heme/10^9^ parasites.

In summary, endocytic null parasites presented a minimal growth defect with both heme source (hemin or Hb) compared to control lines. These parasites presented a protein expression pattern of *Tc*HRG comparable to Dm28c WT epimastigotes under similar treatments. Additionally, IHC of endocytic null parasites was comparable to the WT and complemented line in all conditions evaluated here.

### Overexpression of *Tc*HRG contributes to heme transport in Hb-supplemented parasites

To confirm the role of *Tc*HRG in the uptake of heme derived from Hb, we analyzed the IHC of epimastigotes overexpressing r*Tc*HRG.His-GFP (r*Tc*HRG) when supplemented with Hb as a heme source. Epimastigotes transfected with p*Tc*IndexGW.*Tc*HRG.His-GFP or p*Tc*Index (as a control) were cultured for 48 h in media supplemented with 5 μM heme as hemin or as Hb. Figure 3A shows that epimastigotes expressing r*Tc*HRG incubated with Hb presented an IHC significantly higher compared to control parasites, 2.2 ± 0.1 vs. 1.2 ± 0.1 nmol heme/10^9^ parasites, respectively. Also, the presence of r*Tc*HRG was verified by Western blotting using polyclonal anti-GFP antibodies, as shown in Fig. 3B. Epimastigotes overexpressing r*Tc*HRG incubated with Hb exhibited an incremental increase in IHC analogous to what was reported using hemin as heme source (see Fig. 3A here and [11]).

**Figure 3:**
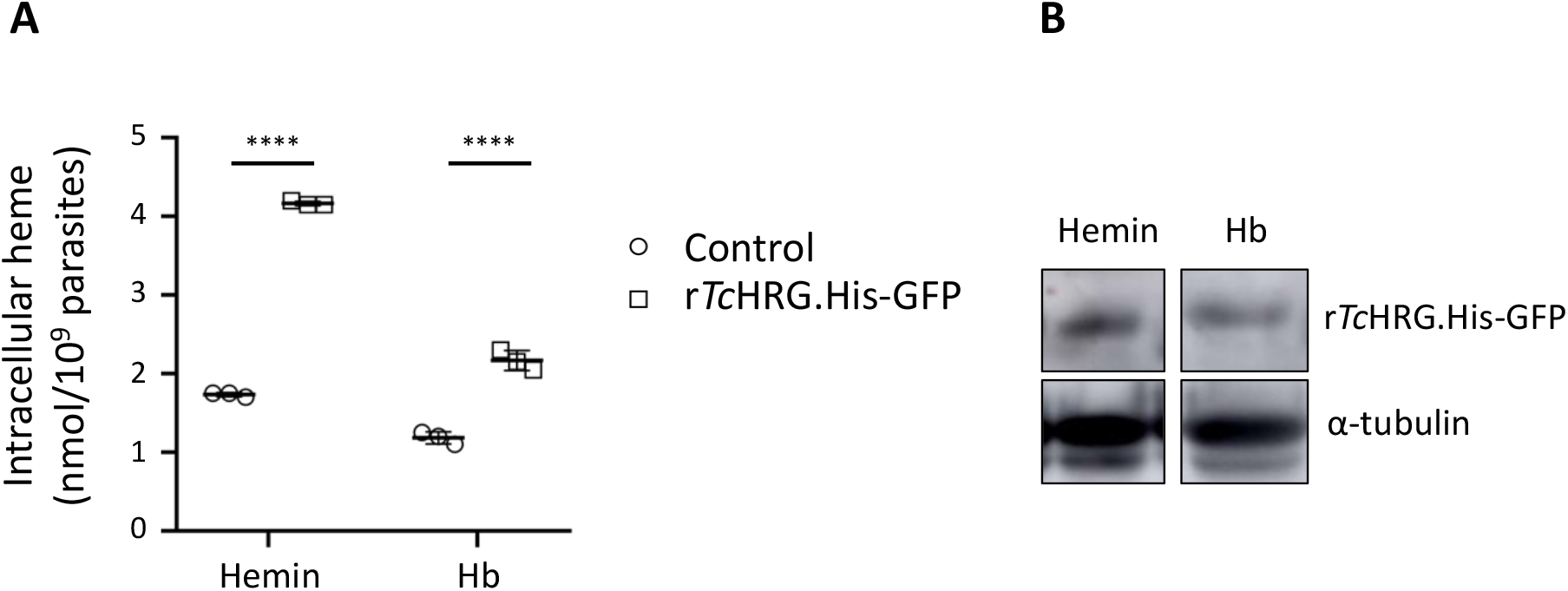
Overexpression of *Tc*HRG leads to an increment on IHC in Hb-supplemented parasites. Control epimastigotes (pLEW 13 p*Tc*Index) and r*Tc*HRG.His-GFP expressing epimastigotes (pLEW 13 p*Tc*Index.GW.*Tc*HRG.His-GFP) (Dm28c strain) were cultured in LIT-10% FBS + 5 µM hemin, then challenged to heme starvation for 48 h and finally transferred to media supplemented with 5 µM hemin or 1.25 µM Hb for 48 h. (A) IHC determined by pyridine method. Data are presented as mean ± SD of 3 independent assays. Statistical significance was determined by two-way ANOVA followed by Sidak’s multiple comparisons test. (****, p < 0.0001; ***, p < 0.001). (B) Western blot assay using anti-GFP antibodies to detect recombinant *Tc*HRG.His-GFP protein in total extracts of r*Tc*HRG.His-GFP overexpressing epimastigotes incubated for 48 h in the conditions mentioned above. Detection of α-tubulin was used as a loading control.

### Endogenous *Tc*HRG has a dual localization

r*Tc*HRG.His-GFP was localized in the FP region of recombinant epimastigotes [11], but the localization of endogenous *Tc*HRG remained elusive. To address the later, first we performed immunofluorescence assays to detect r*Tc*HRG in r*Tc*HRG.His-GFP overexpressing parasites by super-resolution light microscopy using polyclonal a-*Tc*HRG antibodies as primary antibodies. Figure 4A shows that the signal corresponding to anti-*Tc*HRG antibodies (red) overlapped with the green signal of r*Tc*HRG.His-GFP, confirming that these antibodies specifically label *Tc*HRG. Then, we used the same strategy to analyze the localization of endogenous *Tc*HRG in WT parasites incubated with 5 µM hemin or 1.25 µM Hb for 48 h. Additionally, we took advantage of concanavalin A to stain the parasite surface and the cytostome entrance[18]. As shown in Figure 4B, a principal band-shaped signal was observed near the kinetoplast, consistent with a localization in FP region. Additionally, punctual signals were found throughout the cytoplasm, the pattern of these signals suggested that endogenous protein could be localized (at least partially) to the mitochondrion. To corroborate this, we used MitoTracker to stain the parasite mitochondrion as shown in Fig 4C. The green signal of *Tc*HRG partially overlapped with the red signal of MitoTracker, suggesting that endogenous *Tc*HRG could be localized also to the mitochondrial membrane or close to it. On the other hand, we did not observe any difference in the signal patterns of endogenous *Tc*HRG between Y and Dm28c strains nor the incubation conditions (Dm28c not shown) (Fig 4B). In summary, as schematized in Figure 4D, endogenous *Tc*HRG was found in the proximity of the kinetoplast, which is compatible with the position of FP, and partially overlapping with the mitochondrion, while r*Tc*HRG.His-GFP was observed mainly in the FP region.

**Figure 4:**
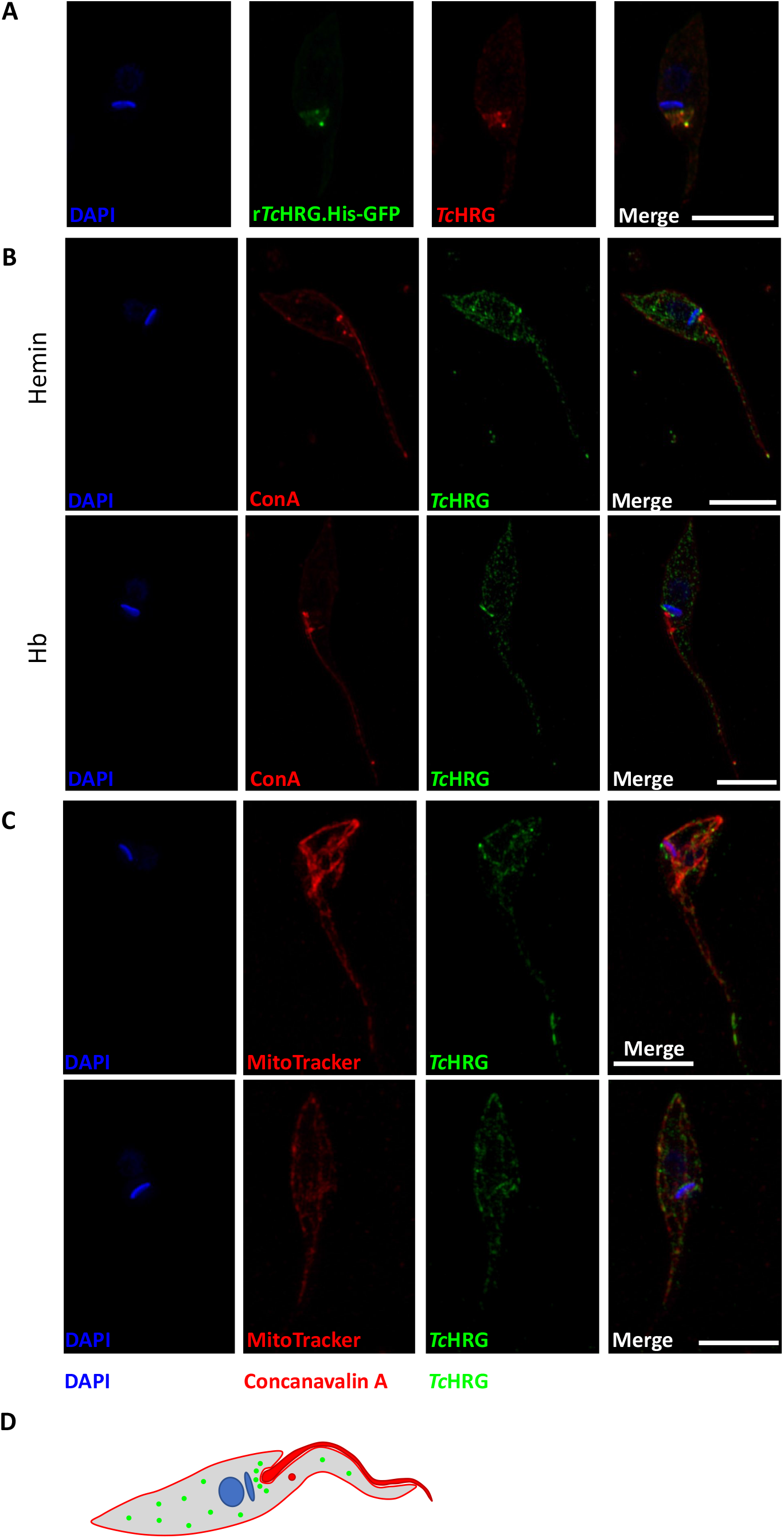
Localization of endogenous *Tc*HRG. Superresolution structured illumination (SR-SIM) microscopy of: (A) Epimastigotes that express r*Tc*HRG.His-GFP (Dm28c strain) cultured in LIT-10% FBS + 5 µM hemin. Polyclonal anti-*Tc*HRG antibodies (red) were used to validate specific union to r*Tc*HRG.His-GFP protein (green). The colocalization analysis (red:green) was performed (Pearson’s coefficient: 0.821 and Manders’ coefficients: 0.988 and 0.969). (B) WT epimastigotes (Y strain) cultured in media supplemented with 5 µM hemin (upper panel) or 1.25 µM Hb (lower panel). anti-*Tc*HRG antibodies were used as to recognize endogenous *Tc*HRG protein (green). Labeling with concanavalin A-rhodamine (ConA, red) was used to identify the parasite’s surface. No difference was observed between Y and Dm28c strain. (C) WT epimastigotes (Y strain) cultured in medium supplemented with 5 µM hemin. a-*Tc*HRG antibodies were used as to recognize endogenous *Tc*HRG protein (green) and MitoTracker™ Red CMXRos were used to label the mitochondria (red). The colocalization analysis (red:green) was performed. Pearson’s coefficient: 0.545 and Manders’ coefficients 0.801 and 0.988 (Upper panel). Pearson’s coefficient: 0.647 and Manders’ coefficients 0.744 and 0.996 (Lower panel). (D) Scheme of *Tc*HRG localization in WT epimastigotes. Nucleus and kinetoplast are schematized in blue, the flagellum and the cytostome entrance are depicted in red. *Tc*HRG is schematized as green dots throughout the parasite’s body and concentrated in the FP area. Nuclei and kinetoplasts in all fluorescent images were stained with DAPI (4,6-diamidino-2-phenylindole, blue). Scale bars: 5 µm.

## DISCUSSION

It was proposed that Hb is the main source of heme *in vivo* for epimastigotes given that it can be obtained from erythrocyte lysis after a bloodmeal in the midgut of the triatomine vector. Then, to obtain heme from Hb, epimastigotes were thought to internalize protein bound heme *via* SPC, degrade it, and then utilize free-heme and peptides. However, conversely to *T. brucei* and *Leishmania spp*., no specific receptor that mediates Hb endocytosis or putative ORF that may fulfill this function has been described or found in *T. cruzi* genome and very little is known about the endocytic process through the SPC. On the other hand, Hb can be degraded via external peptidases to release free heme and peptides, although triatomines present mechanisms to rapidly get rid of free heme to avoid oxidative damage [19]. In this work, we present a model for Hb-derived heme uptake, explaining the role of *Tc*HRG in this process.

The growth performance of epimastigotes was not affected when Hb was used as heme source in axenic culture, moreover, higher concentrations of Hb did not cause the same negative effect on growth as is observed when equivalent concentration of hemin was added [12]. This phenomenon may be explained because both sources are incorporated via different pathways or kinetics allowing the parasite to distribute and correctly store the cofactor, and/or because heme embedded within Hb produces less oxidative damage to the lipids and proteins of the plasma membrane as compared to free heme[20]. Additionally, the concentration of intracellular heme differed between both heme sources during the first 48 h after they were added to the medium, but reached similar values at 96 h, supporting the idea that time is needed to degrade extracellular hemoglobin and thus release Hb-derived heme for uptake and intracellular utilization by epimastigotes.

*Tc*HRG (formerly *Tc*HTE) is an essential player in the control and regulation of free heme uptake, as its amount (both mRNA and protein) are adjusted according to the intracellular heme status[12]. Our results have shown that the *Tc*HRG protein signal decreases when both heme sources are used; however, it is still detected 24 h – 48 h post Hb addition. Furthermore, mRNA levels dropped about 25 % with Hb whereas about 50 % with hemin, in both cases when the heme source was added after heme starvation. This behavior agrees with the observed intracellular heme levels and strongly suggest that, despite that how Hb-derived heme enters the cell, *Tc*HRG plays a crucial role sensing intracellular heme and responding to it.

As mentioned before, Hb might be internalized via SPC however, the growth performance of endocytic null epimastigotes (Δ*Tc*Act2) was similar in the presence of both heme sources, although the final parasite number was slightly lower as compared to WT and complemented (*Tc*Act2.Ty) lines in the same conditions. This minor phenomenon is possibly due to the inability to obtain some additional minor nutrients by endocytosis. Regardless, when the intracellular heme concentration was measured in WT, Δ*Tc*Act2 and *Tc*Act2.Ty they all demonstrated a similar profile. Heme starvation caused a drop in the intracellular levels, and, although IHC was lower when Hb was used as heme source compared to hemin, the same behavior is observed in Dm28c epimastigotes. Independently of the heme source supplemented, endocytic null parasites exhibit similar growth with both and can reach WT intracellular heme levels, indicating that epimastigotes rely on mechanisms independent of endocytosis to take up heme from the environment. Importantly, the *Tc*HRG protein is present at the same level in all three lines, being almost indetectable when hemin was used and still detected 48 h after the addition of Hb, confirming the role of this protein in control of heme homeostasis independently of the heme source. Additionally, the overexpression of r*Tc*HRG led to an increase in the intracellular heme content in parasites supplemented with Hb comparable to the effect observed when hemin is used as free heme source [11]. These data suggest that the incorporation of free heme and heme derived from Hb could proceed by same mechanism but with different timing. One possible explanation for this phenomenon is that the parasite may contribute (at least partially) to extracellular Hb breakdown *via* secreted [21] or surface proteases [22], then “free” heme derived from Hb degradation required extra time to be incorporated compared to the addition of hemin to the medium. The hypothesis that parasite proteases might contribute to protein digestion in the lumen of the triatomine midgut was suggested by García and colleagues, although the details of this process remain poorly understood [23].

Taking advantage of super-resolution microscopy, it was possible to observe recombinant as well as endogenous *Tc*HRG using anti-*Tc*HRG antibodies. r*Tc*HRG was detected in the proximity of the FP as previously reported [11], meanwhile the endogenous protein signal was visualized for the first time, and it was localized next to the FP (close to the kinetoplast) and as multiple puncta in the cytoplasm. The localization of r*Tc*HRG signal only in FP region could be a consequence of the structure of the fusion protein where, probably, the presence of the GFP moiety would prevent the migration of the protein to other cellular regions. Interestingly, the intracellular signal of *Tc*HRG did not change according to the heme source. Its localization in the FP region supports that *Tc*HRG plays a role in both heme uptake and homeostasis control, in agreement with the hypothesis that permeases and transporters in trypanosomatids also play roles in sensing [24]. The intracellular signals of *Tc*HRG exhibited partial superposition with mitochondrial signal, suggesting a more complex role for this protein; probably involved in intracellular heme trafficking or sensing. This localization differs from the observed in other trypanosomatids where HRGs were observed in intracellular vesicles [6,9], but in all cases it was suggested a role in heme trafficking. One possible role of mitochondrial *Tc*HRG could be the sensing of intracellular heme status and/or intracellular heme trafficking in this organelle. Results presented in this work indicate that the main heme entrance pathway in *T. cruzi* epimastigotes is the same for both heme sources (hemin and Hb) and it is mediated by *Tc*HRG. We explain how epimastigotes control heme homeostasis regulated by *Tc*HRG when Hb is available as a heme source, as summarized in Figure 5. Hb can be externally degraded by surface or secreted proteases produced by either the parasite or the insect vector [23], and free heme uptake is enhanced by *Tc*HRG at the FP region. Although not examined in this study, Hb might be endocytosed *via* SPC. More importantly, independent of the heme source, epimastigotes sense intracellular heme status presumably *via* intracellular *Tc*HRG. Once the optimal concentration range is reached, a still unknown signal may trigger *Tc*HRG to turn off to avoid further heme uptake which is likely toxic, as was previously reported [12].

**Figure 5:**
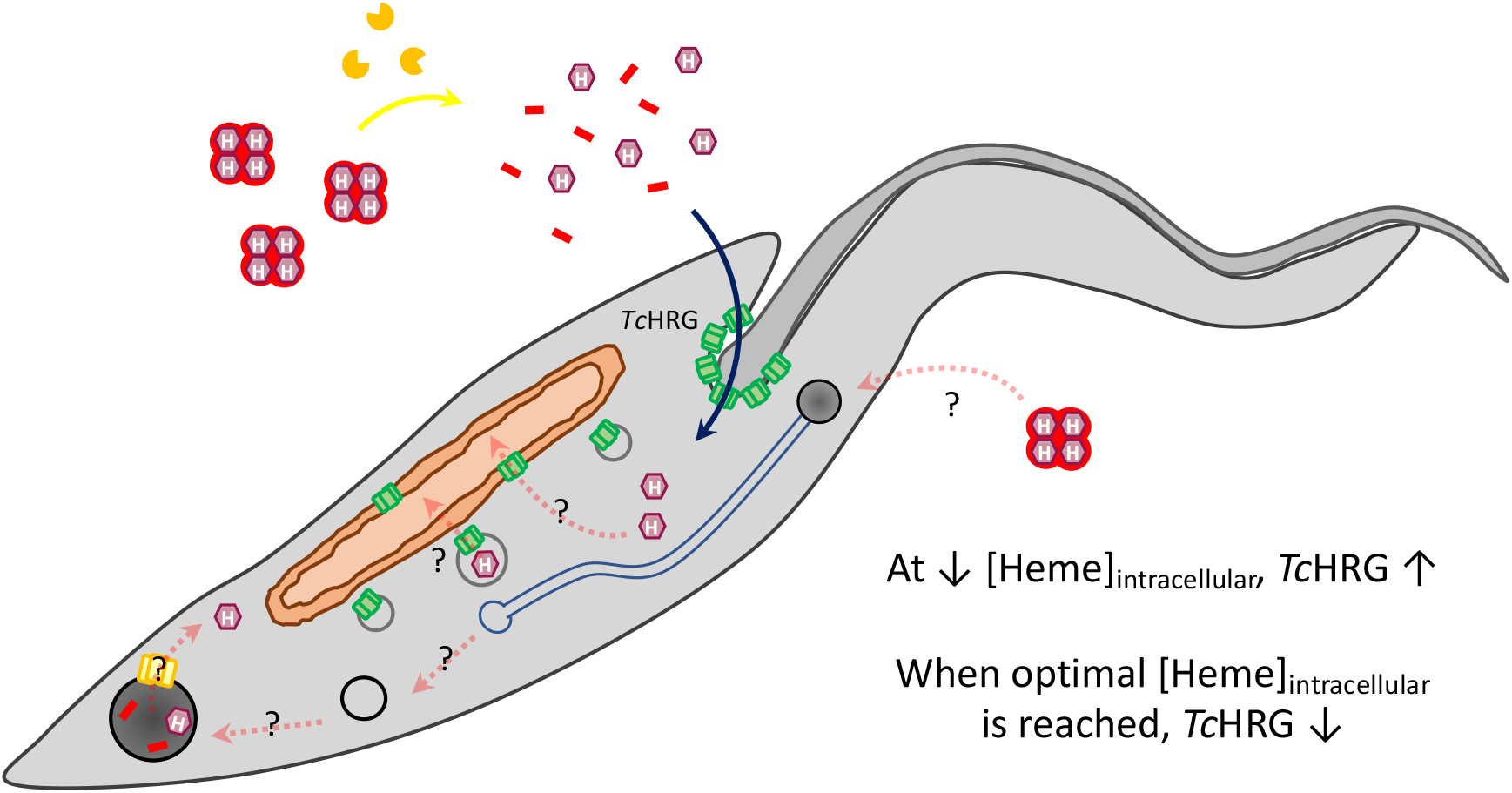
Scheme of the proposed model of heme uptake in epimastigotes of *Trypanosoma cruzi*. When the epimastigote senses that the IHC is below the optimal range, *Tc*HRG expression increases, promoting the incorporation of heme derived from externally degraded Hb in the FP region. Heme should be then distributed and incorporated into hemoproteins, and mitochondrial *Tc*HRG could also be involved in this process promoting heme transport to the mitochondria. Hb might also be endocytosed through the SPC and internally processed to obtain free heme, that may be exported to the cytosol or stored in reservosomes. Once the parasite obtains enough heme to satisfy its nutritional requirements, likely *via* intracellular *Tc*HRG, its expression is downregulated to maintain heme homeostasis.

In summary, we present here an extended model to explain Hb-derived heme uptake in epimastigotes of *T. cruzi*. Our results reinforce the relevance of *Tc*HRG in the crucial process of heme transport and homeostasis independently of the source, constituting a key player in *Trypanosoma cruzi*. For these reasons, elucidation of the complete heme uptake pathways will contribute to the identification of other novel essential proteins and generate new strategies for Chagas’ disease treatment.

### Experimental procedures

#### Reagents

Fetal Bovine Serum (FBS) (Internegocios SA) was heat-inactivated at 56°C for an hour prior to use. Hemin (Frontier Scientific) solution stock was prepared as previously described (Merli 2016). Heme concentration in hemin stock solution was confirmed by spectroscopic measurements at 385 nm, ε^385^= 58400 M^-1^cm^-1^ [25]. Lyophilized bovine Hb (Sigma) stock solution was prepared in PBS to a final concentration of 0.1 mM, sterilized by filtration using a 0.22 μm syringe disposable filter and stored at -80°C. Concentration in Hb stock was confirmed by indirectly measuring heme by basic piridine method described below, considering that one Hb molecule contains four heme molecules. Integrity of Hb secondary structure was verified by circular dichroism spectroscopy and SDS-PAGE.

#### Parasites

*T. cruzi* Dm28c strain was used to analyze parasite growth, expression of endogenous *Tc*HRG (WT) and to study the effect of r*Tc*HRG overexpression and localization in medium supplemented with Hb (pLEW13 p*Tc*Index and pLEW13 p*Tc*Index.GW.*Tc*HRG.6His-GFP lines). Analysis of endocytosis suppression was performed using *T. cruzi* Y strain (WT, Δ*Tc*Act2, and *Tc*Act2.Ty lines).

Epimastigotes were routinely maintained in mid-log phase by periodic dilutions in Liver Infusion Tryptose (LIT) medium supplemented with 10% heat inactivated FBS (LIT-10% FBS) and 5 μM hemin (Pagura 2020), at 28 °C. Prior to each experiment described in this work, epimastigotes were collected, washed with PBS and transferred to LIT-10% FBS without heme source added for 48 h (“heme starvation”).

#### Growth curves

Dm28c WT epimastigotes routinely maintained LIT-10% FBS + 5 μM hemin were challenged to heme starvation for 48 h. Then, parasites were collected by centrifugation at 3500 g for 5’, washed with PBS and transferred to LIT-10% FBS without heme (0 µM) or supplemented with 5, 20, or 50 µM hemin or 1.25, 5, and 12.5 µM Hb (equivalent to 5, 20, or 50 µM heme as Hb). The number of cells was monitored daily for 14 days. On day 7, cultures were diluted to the parasite concentration measured on day 1 and the growth curve was followed for another week. On the first-, third-, seventh-, and fourteenth-days parasites samples were collected and prepared for Western Blot analysis as described below and the morphology was verified by optical microscopy. Similarly, Y epimastigotes (WT, Δ*Tc*Act2, and *Tc*Act2.Ty lines) were heme starved for 48 h and transferred to LIT-10% FBS without (0 µM) or supplemented with 5 µM hemin or 1.25 µM Hb. The growth curve was followed for one week. The results are expressed as mean ± SD of three independent experiments. Cell growth was monitored by cell counting using Wiener lab. Counter 19 Auto Hematology Analyzer (Wiener Laboratorios SAIC, Rosario, Argentina) configured for parasite number measurements and Neubauer chamber.

#### Western blot

Total protein from cell-free extracts were obtained and processed as described previously [12] with minor modifications. 5 × 10^6^ cells/well were resolved by electrophoresis on a 12% SDS-polyacrylamide gel. *Tc*HRG detection was performed with rabbit polyclonal anti-*Tc*HRG antibodies (1:10000). r*Tc*HRG.6His-GFP expression was corroborated using anti-GFP antibodies (1:1000) (Santa Cruz Biotechnology, Inc.). In both cases, peroxidase-labeled anti-rabbit IgG (1:30000) (Calbiochem) were used as secondary antibodies. Loading control was performed with anti-α-tubulin clone TAT-1 antibodies (a gift from K. Gull, University of Oxford, U.K.), using peroxidase-labeled anti-mouse IgG antibodies (1:5000) (GE Heathcare) as secondary antibodies. Bound antibodies were detected with ECL Prime Western Blotting Detection kit (GE Healthcare). Anti-*Tc*HRG antibodies were generated as described previously [12].

### RNA isolation, reverse transcription PCR (RT-PCR) and quantitative real-time PCR (qRT-PCR)

After heme starvation, Dm28c WT epimastigotes were collected, washed with PBS and challenged to grow in LIT-10% FBS without (0 µM) or supplemented with 5 µM hemin or 1.25 µM Hb. Samples in triplicates were collected for RNA isolation prior transferring parasites to the different conditions (t_0_) and after 24 hours of incubation (t_24_). Total mRNA isolation, treatment and quantification was carried out as described previously (Pagura 2020). cDNAs were synthesized through a RT reaction (M-MuLV, Thermo-Scientific) using 0.5 µg of total RNA. cDNA samples were used as template for Quantitative Real-time PCR performed in an Applied Biosystems StepOne™ Real-Time PCR System Thermal Cycling Block using the EvaGreen fluorescence quantification system (Solis BioDyne). qRT-PCR reaction was conducted as previously described (Pagura 2020). The results are expressed as mean ± SD of three technical replica from one representative of two independent experiments (biological replica).

#### Immunofluorescence assays

Epimastigotes (Y and Dm28c strains) are routinely grown in LIT-10% FBS + 5 μM hemin were subjected to heme starvation for 48 h. Then, parasites were harvested, washed with PBS and transferred to LIT-10% FBS without (0 µM) or supplemented with 5 µM hemin or 1.25 µM Hb for 48 h.

Mounted samples of epimastigotes for fluorescence imaging were prepared and labeled with 10 µg/ml rhodamine-conjugated concanavalin A (Vector Laboratories) as described previously (Chasen 2019). Polyclonal anti-*Tc*HRG antibodies were used as primary antibodies in WT and r*Tc*HRG.His-GFP overexpressing parasites. As secondary antibodies, goat anti-rabbit IgG Alexa Fluor 488 (Thermo Fisher) were used in WT epimastigotes, while goat anti-rabbit IgG Alexa Fluor 568 (Thermo Fisher) were used in r*Tc*HRG.His-GFP overexpressing parasites. MitoTracker™ Red CMXRos (Thermo Fisher) was used to dye epimastigotes mitochondria.

All the images were acquired with Zeiss Elyra S1 structured illumination microscope (Center for Tropical and Emerging Diseases Biomedical Microscopy Core, Athens, GA) and were processed using the ImageJ software.

#### Heme content analysis

After heme starvation, parasites were harvested, washed twice with PBS and transferred to the corresponding medium (supplemented with 5 µM hemin, 1.25 µM Hb, or without heme) for 48 h or 96 h. Epimastigotes were harvested and washed three times with PBS. Each determination required 150 × 10^6^ epimastigotes. Intracellular heme content of epimastigotes was determined by basic pyridine method described in Berry *et al*. [26], adapted in our laboratory to perform measurements in epimastigotes samples, as we described previously [27]. The results are expressed as mean ± SD of three independent experiments (biological replicates), each containing two technical replicates.

#### Statistical analysis

All the assays were independently reproduced at least 2–3 times. Statistically significant differences between groups were analyzed using GRAPHPAD PRISM version 6.00 for Windows (GraphPad Software, San Diego, CA), as described in each experiment.

## Acknowledgments

We are grateful to the government of the province of Santa Fe for awarding ET the “Mobility Scholarship with a Gender Perspective” (2022).

## Author contributions

J.A.C. and E.T. conceived, designed and supervised the project, E.T. performed most of designed experiments, C.B.D.C. performed qPCR assay, N.M.C obtained the microscopy images. J.A.C., E.T. and R.D.E. discussed the results. J.A.C. and E.T. wrote the manuscript with contributions from all other authors.

## Abbreviations

ANOVA: Analysis of variance
ConA: concanavalin A
FBS: Fetal Bovine Serum
FP: flagellar pocket
Hb: Hemoglobin
HRG: Heme Responsive Gene
IHC: Intracellular heme content
LIT: Liver Infusion Tryptose
PBS: Phosphate buffered saline
qRT-PCR: quantitative real-time PCR
r*Tc*HRG: recombinant *Tc*HRG
SPC: Citostome-cytopharinx complex
*Tc*HRG: Trypanosoma cruzi Heme Response Gene
WT: Wild Type

